# Decoding explicit and implicit representations of health and taste attributes of foods in the human brain

**DOI:** 10.1101/2021.05.16.444383

**Authors:** Elektra Schubert, Daniel Rosenblatt, Djamila Eliby, Yoshihisa Kashima, Hinze Hogendoorn, Stefan Bode

## Abstract

Obesity has become a significant problem word-wide and is strongly linked to poor food choices. Even in healthy individuals, taste perceptions often drive dietary decisions more strongly than healthiness. This study tested whether health and taste representations can be directly decoded from brain activity, both when explicitly considered, and when implicitly processed for decision-making. We used multivariate support vector regression for event-related potentials (as measured by the electroencephalogram) occurring in the first second of food cue processing to predict ratings of tastiness and healthiness. In Experiment 1, 37 healthy participants viewed images of various foods and explicitly rated their tastiness and healthiness, whereas in Experiment 2, 89 healthy participants indicated their desire to consume snack foods, with no explicit instruction to consider tastiness or healthiness. In Experiment 1 both attributes could be decoded, with taste information being available earlier than health. In Experiment 2, both dimensions were also decodable, and their significant decoding preceded the decoding of decisions (i.e., desire to consume the food). However, in Experiment 2, health representations were decodable earlier than taste representations. These results suggest that health information is activated in the brain during the early stages of dietary decisions, which is promising for designing obesity interventions aimed at quickly activating health awareness.

---

Obesity is a global epidemic, and excess body weight is associated with increased risk of chronic diseases, estimated to cause nearly three million annual deaths worldwide (Lauby-Secretan et al., 2016; Stevens et al., 2012). While obesity is a multifaceted issue, there is a strong link to poor food choices leading to a diet high in sugar and fat (Fock & Khoo, 2013). During dietary decisions, although people tend to consider both tastiness and healthiness of foods (Sullivan et al., 2015; Hare et al., 2009), these attributes are often in conflict, and foods considered unhealthy tend to have higher perceived tastiness (Raghunathan et al., 2006; Mai & Hoffmann, 2015). This is problematic because taste attributes tend to drive dietary decisions more strongly (Tepper & Trail, 1998; Pollard et al., 1998; Markovina et al., 2015). To inform the development of effective interventions against obesity, it is therefore important to understand how both perceptions are processed in the brain.

Compared to health, taste is a less abstract, lower-level representation, thought to be processed and integrated into decisions quickly and automatically (Sullivan et al., 2015; Liberman & Trope, 2008; Motoki et al., 2018). Supporting this, several studies have used electroencephalography (EEG) to suggest that the tastiness of foods is reflected by event-related potential (ERP) components occurring during the first second of viewing a food stimulus (Rosenblatt et al., 2018; Schwab et al., 2017; Sarlo et al., 2013; Schubert et al., 2021). In particular, the late positive potential (LPP), occurring approximately 450-1000 ms after the presentation of a stimulus and previously implicated in emotional responding and arousal (Schupp et al., 2000; Pastor et al., 2008; Gable & Harmon-Jones, 2010), is thought to reflect the caloric or appetitive value of stimuli (Rosenblatt et al., 2018; Schwab et al., 2017; Schubert et al., 2021). Images of high-calorie foods, including meat dishes and sweets, have been shown to elicit larger LPP amplitudes than vegetable images (Schwab et al., 2017). There is also evidence that the LPP reflects perceived tastiness, with higher amplitudes found in response to images of sugar-sweetened beverages perceived to be tasty compared to non-tasty (Schubert et al., 2021). This study also found differences between these two categories in the N1 component, which occurs approximately 100 ms after the onset of a stimulus and has been implicated in early attentional processing (Schubert et al., 2021; Carbine et al., 2018).

These studies suggest that when people view food cues, taste perceptions may be activated quickly and automatically, during the first second of processing. However, it is unclear whether this also applies to health perceptions. There is evidence that perceptions of health attributes are activated and represented in the ventromedial prefrontal cortex during dietary decisions (Cosme et al., 2020); however, it is unclear whether this occurs as quickly as the activation of taste attributes. Preliminary evidence from a study using computer mouse trajectory tracking suggests that health attributes are processed and integrated into dietary decisions approximately 195 ms later than taste attributes (Sullivan et al., 2015). It is hence possible that neural representations of healthiness are formed later than taste representations, which may contribute toward people’s tendency to choose to eat tasty but unhealthy foods (Tepper & Trail, 1998; Pollard et al., 1998; Markovina et al., 2015). However, no studies to date have been able to directly investigate the activation of neural health and taste representations at a suitable temporal scale.

## The Present Study

Previous ERP studies (Schwab et al., 2017; Schubert et al., 2021) hint at potential processing differences between high- and low-calorie foods in time. Our study, however, went beyond that and investigated whether and when taste and health attributes of foods were neurally represented in brain activity on a trial-by-trial basis. For this, we used a novel multivariate regression approach, which involves training a classifier to predict ratings for healthiness and tastiness based on spatially distributed patterns of neural activity in small time windows, and then testing its performance on new data. Because it utilises the entire distributed EEG signal (Bode et al., 2019), it becomes possible to “read out” cognitive information, even abstract information such as semantic attributes of objects (Bode et al., 2014) and subtle changes in emotional states (Schubert et al., 2020), which has not been achievable in the past.

In the present study, we aimed to determine whether taste and health perceptions are decodable in the brain during the early stages of food cue processing. In Experiment 1, we examined representations of these attributes during explicit evaluation of tastiness and healthiness. This involved predicting both tastiness and healthiness ratings based on neural patterns recorded immediately after presenting a food image, but before a response could be prepared. In Experiment 2, we used a similar approach to examine health and taste representations when individuals made consumption decisions about food items, but were not explicitly told to consider tastiness and healthiness. For this second experiment, we reanalysed data from a previous study (Rosenblatt et al., 2018). We hypothesised that health and taste attributes would be represented in brain activity in both studies. We further hypothesised that evidence for both health and taste decoding would be found in Experiment 2 before the consumption decision outcome was represented, which could be interpreted as evidence that participants integrated both sources of information to inform their decisions.

## Experiment 1

### 2.1 Method

#### 2.1.1 Participants

The sample comprised 39 participants, all right-handed, fluent in written and spoken English, having normal or corrected-to-normal vision, and no dietary restrictions or eating disorder history. As there is currently no literature on calculating statistical power for multivariate EEG analyses, the sample size was based on previous studies using EEG to examine neural responses to food cues (e.g., Meule et al., 2013; Sarlo et al., 2013). Participants were asked to fast for four hours before the experiment to control for hunger levels. Two participants were excluded because their results fluctuated more than ±3 SD between analysis time windows throughout the entire trial, indicating a failure to fit the regression model (see below). In the final sample, there were 37 participants (age range = 18 to 36 years; *M* = 24.08 years, *SD* = 4.74; 29 females, 8 males). Before the task, participants gave written informed consent, and afterwards they were debriefed and given a voucher payment of AUD$20. The experiment was approved by the Human Research Ethics Committee (ID1955772) of the University of Melbourne and conducted in accordance with the Declaration of Helsinki.

#### 2.1.2 Stimuli

Stimuli were 174 images of foods from the Food-Pics database (450 x 338 pixels), selected based on normative palatability ratings and food categorisations from omnivorous participants (Blechert et al., 2019). The stimuli represented a variety of food types (fruit, vegetables, chocolate, fish, meat, nuts, snacks/meals) and a wide range of subjective palatability.

#### 2.1.3 Questionnaires

Participants completed several questionnaires before the experiment, including demographic information about age, gender, level of education, height, and weight. They then rated their hunger level on a scale of 1 (not at all) to 9 (extremely hungry) and indicated how many hours had passed since their last consumed snack and meal. Finally, they completed several questionnaires about eating behaviour (see Supplementary Material 1).

#### 2.1.4 Procedure

Each trial began with a fixation cross (1.5 s), followed by a food image (2 s; Figure 1). Then, while the food image remained onscreen, a question appeared underneath – either “How much do you enjoy the taste of this food?” or “How healthy do you consider this food to be?”. Participants answered using a continuous sliding scale, ranging from “Not at all” to “Very much”. The underlying values of 0 to 100 were not visible to participants. The ends of the scale were randomly reversed on half of the trials to prevent motor preparation in the initial image presentation period. Participants had 10 seconds to answer by moving the slider to the desired position and clicking the mouse. If no response was given after 10 seconds, the current position of the slider was recorded as a response. Each image was shown twice, once with each question. The experiment consisted of 348 trials, split into twelve experimental blocks. In each block, only one of the questions was asked, alternating between blocks. The question displayed in the first block was randomised across participants. The whole task took approximately one hour, including EEG setup and pack-up.

**Figure 1.**
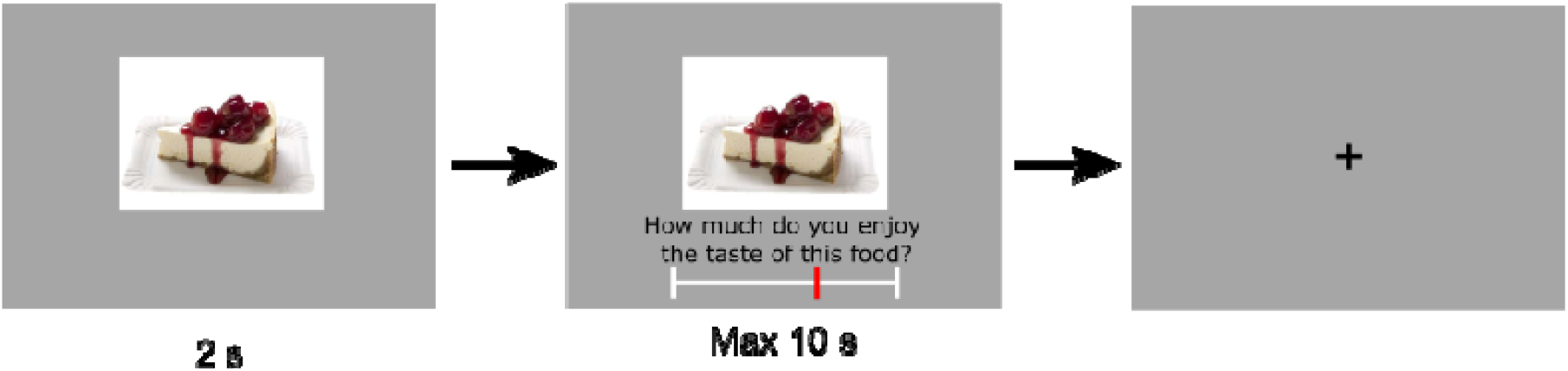
Trial structure for Experiment 1. Participants viewed each image for 2 s before a continuous slider scale appeared, asking them to rate either taste or health, depending on the current block. Health and taste rating blocks were alternated, and all 174 images were rated on each attribute once.

#### 2.1.5 EEG Recording & Preprocessing

Electrophysiological activity was recorded using a 64-channel BioSemi Active II system at a sample rate of 512 Hz, and recording bandwidth of DC-102 Hz. Ag/AgC1 electrodes were attached to a fabric cap according to the International 10-20 system, with additional electrodes beside and below the left eye (horizontal and vertical electrooculogram), and above the left and right mastoids (reference electrodes). Electrode offsets were kept within ±50 μV. Using EEGLab v14.1.2 (Delorme & Makeig, 2004), the data were firstly re-referenced to the average of the mastoids, then high-pass (0.1 Hz) and low-pass (30 Hz) filtered (EEGLab FIR Filter New, default settings). They were segmented into epochs beginning 100 ms before an image was presented (used as the baseline), and ending 1,200 ms after image onset, to capture the first period of visual and semantic processing while avoiding time periods that already contained activity related to motor preparation. Epochs containing muscle and skin potential artefacts were identified via visual inspection and removed. Excessively noisy channels were removed and interpolated using spherical spline interpolation (this occurred for 23 participants; average 2.56 channels interpolated per participant). An independent components analysis (ICA) as implemented in EEGlab was used to identify and remove eye movements, saccades, and blinks. If amplitudes at any channel exceeded ±150 μV, the epochs were excluded from analyses.

#### 2.1.6 Decoding Analysis

For the multivariate pattern analyses (MVPA), linear support vector regression (SVR) as implemented in the Decision Decoding Toolbox v1.0.5 (DDTBOX; Bode et al., 2019) was used to examine the association between brain activity and continuous taste or health ratings for each trial. For each participant, two analyses were performed – one using taste ratings and brain activity recorded after image presentation in taste blocks, and the other using health ratings and brain activity recorded after image presentation in health blocks. A “sliding window” 20 ms wide moved through each epoch of the EEG data in overlapping 10 ms steps. Within each window, all ten data points from the 64 EEG channels were transformed into vectors of spatial brain activity patterns (10 x 64 = 640 features in total) associated with each trial. These vectors were then used to predict the respective ratings. In this process, a linear SVR model (standard cost parameter C = 0.1, interfacing LIMSVM; Chang & Lin, 2011) was trained on a randomly selected 90% of the data, then tested on the remaining 10%. This process was repeated independently using a 10-fold cross-validation procedure, until all data sets had been used as test data once while training on all other sets. The cross-validation process itself was then repeated ten times with newly drawn random data, to obtain a conservative estimate of decoding performance based on average classifier performance from all 10×10 iterations, circumventing drawing biases.

Each analysis time window produced an average Fisher-Z-transformed correlation coefficient between predicted and true rating scores. To generate an empirical chance distribution for statistical testing, this procedure was repeated with the same data but with the ratings randomly shuffled across trials (c.f., Bode et al., 2019; Schubert et al., 2021). Finally, corrections for multiple comparisons were made using cluster-based permutation tests based on the cluster mass statistic, which takes advantage of the statistical non-independence of the adjacent analysis time windows. Note however, that these corrections might be overly conservative given that for each time window, an independent, empirical chance distribution was generated for testing. We therefore report both corrected and uncorrected results.

### 2.2 Results

Mean responses were 65.27 (*SE* = 4.64) for taste and 47.66 (*SE* = 5.65) for health. As expected, the correlation between taste and health (Pearson’s *r*) varied substantially between participants, ranging from -.15 to .87 (*mean r* = .10, *SD* = .33). Overall, there was a significant correlation between average taste and health ratings, *r*(35) = .33, *p* = .049.

Taste ratings could be predicted significantly above chance in several time windows after image onset, starting at 530 ms (see Figure 2A). Above chance prediction was also possible for health ratings, starting slightly later at 640 ms after image onset (see Figure 2B). Figure 2 further reports all uncorrected significant time windows. For test statistics at each timepoint, see Supplementary Table S3.

**Figure 2.**
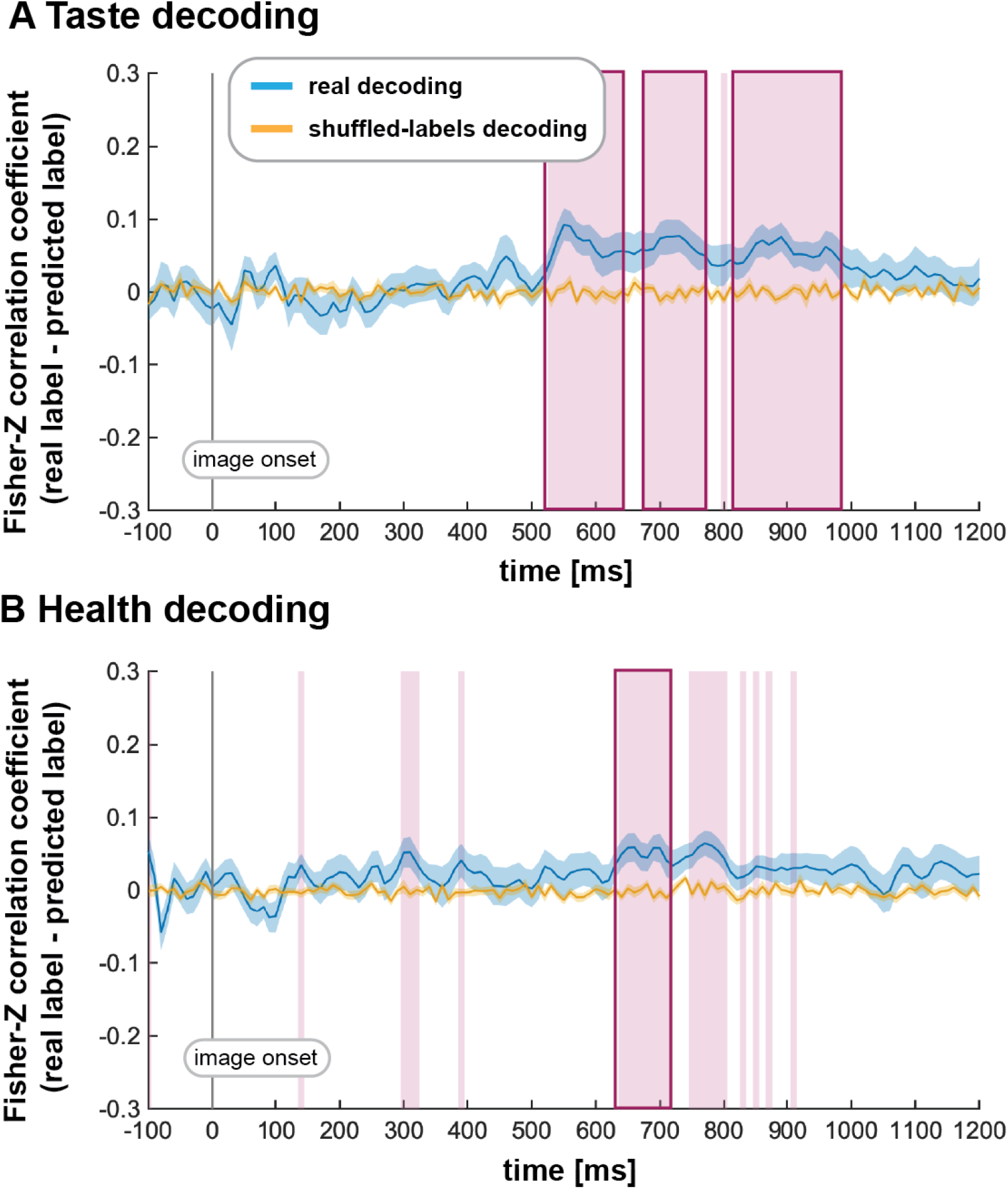
Decoding results for A) taste and B) health ratings. Blue lines represent Fisher-transformed correlation coefficients between predicted and true labels (taste and health ratings), while yellow lines represent decoding results based on shuffled labels (empirical chance distribution). Pink vertical bars highlight time points at which the decoding results significantly differed from the shuffled decoding results (*p* < .05); bolded pink bars represent time windows that remained significant after corrections for multiple comparisons.

## Experiment 2

The results from Experiment 1 show that both taste and health attributes could be decoded from brain activity when participants were explicitly instructed to consider them. In Experiment 2, we examined whether such representations were detectable when participants were not directly prompted to consider either attribute but instead made decisions about how much they wanted to consume foods. For this, we reanalysed a large data set recorded during the first stage of a previous study in which participants made exactly these decisions (Rosenblatt et al., 2018).

### 3.1 Method

#### 3.1.1 Participants

Ninety-six participants were recruited, all right-handed and fluent in English, and free from any history of eating disorders, or health, religious, or ethical circumstances that prevented them from consuming particular snack foods. Participants were asked to fast for four hours before the experiment to control for hunger levels, and were naïve to the purpose of the study. Three participants were removed due to technical problems with data recording, and a further four were removed post hoc due to excessive artifacts in their EEG data. The final sample comprised 89 participants (62 female, 27 male), aged 18 to 50 years (*M* = 22.56, *SD* = 4.78). For the analysis of taste and health, only participants who demonstrated sufficient variability in the relevant outcome variable (taste or health rating SD > 0.7) could be included to achieve a successful regression model fit. As explained below, this experiment assessed ratings on a fixed four-point scale rather than in a continuous fashion as in Experiment 1, which naturally reduced the variability of responses and therefore the number of usable datasets. This selection step resulted in a sample of 55 participants for the taste analysis (39 female, 16 male; *M*_age_ = 22.55 years, *SD* = 4.26) and 85 participants for the health analysis (61 female, 24 male; *M*_age_ = 22.54 years, *SD* = 4.87). Note that using even stricter variance criteria (e.g., SD > 0.8) further reduced the number of data sets and was therefore not included here. Participants gave written informed consent before commencing the task, and were debriefed afterwards and compensated with payment of AUD$20. The experiment was approved by the Human Research Ethics Committee (ID 1443258) of the University of Melbourne and conducted in accordance with the Declaration of Helsinki.

#### 3.1.2 Stimuli

Stimuli were 100 colour images (500⨯ 500 pixels) of snack foods, including nuts, vegetables, fruits, chips, and sweets, selected through another pre-study where a separate sample rated 492 foods on their tastiness, healthiness, and familiarity (see Rosenblatt et al., 2018). 100 food items were then selected based on the average health and taste ratings for the 492 food images, covering the full spectrum of healthiness and tastiness. Note that this set of snack foods differed from the foods in Experiment 1. The original purpose of the study was to investigate the effect of health warning labels (which were introduced in a second part of this study that was not relevant here) on changes in consumption decisions.

#### 3.1.3 Questionnaires

Participants completed a questionnaire recording demographic information, as well as information on smoking status, dieting status, eating disorder history, and food preferences. They also completed a series of questionnaires measuring eating behaviour (see Supplementary Material 2).

#### 3.1.4 Procedure

Before the experiment, participants rated their hunger levels (1 = not at all hungry; 8 = extremely hungry) and recorded the number of hours since their last meal. They then completed a computer-based task during which EEG was recorded, and of which only the first two task stages are relevant here (see Rosenblatt et al., 2018).

In the first stage, participants rated each item on healthiness and tastiness. Each trial began with a fixation point (2 s to 6 s), followed by a food image (3 s). They then had three seconds to respond to one of two possible questions: “Do you like the taste of this food?” or “Do you think this food is healthy?”. Responses were given on a four point scale (1 = Strong No, 2 = Weak No, 3 = Weak Yes, 4 = Strong Yes) by pressing a button on the computer keyboard, with assignment of keys randomised trial-by-trial (note that this differs from Experiment 1). Each food image appeared twice, such that both health and taste ratings were obtained for each image, resulting in 200 trials in total, distributed over four blocks of 50 trials.

Participants then completed the decision stage, which consisted of 50 trials in which half of the food images were presented (the other half was reserved for a second decision stage, after the exposure of health warning messages, that is not relevant here). On each trial, a fixation point was presented (2 s to 6 s), followed by a food image (3 s; see Figure 3). The question “Would you like to eat this food at the end of the experiment?” then appeared. Responses were made on a continuous scale ranging from “Strong No” to “Strong Yes” (midpoint not selectable), using a mouse. The underlying range of -100 to 100 was not visible to participants. The ends of the scale were again randomly reversed trial-by-trial. Participants were informed that their decisions on each trial would directly impact the probability of receiving an item for consumption after the experiment, incentivising realistic choices.

**Figure 3.**
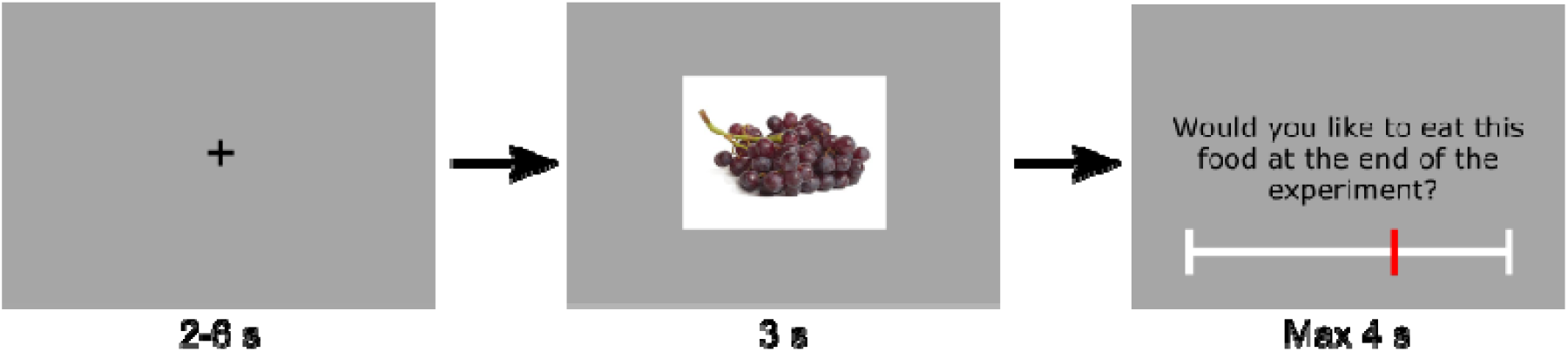
Trial structure for the decision stage in Experiment 2. A fixation point was shown for 2 to 6 s before an image of a food item appeared for 3 s. A continuous slider scale then appeared for 3 s, asking participants how much they would like to eat the food item at the experiment’s conclusion.

Finally, participants completed the questionnaires collecting demographic and eating-related information. At the end of the experiment, participants received one randomly selected food item that they had responded “Yes” to in the decision stages.

#### 3.1.5 EEG Recording & Preprocessing

The EEG recording setup mirrored that of Experiment 1 (see Section 2.1.5), with some small differences - the data in Experiment 2 were filtered online with 70 Hz low-pass filter, which however did not make a difference to the following analyses. The data from the decision stage were then separated into epochs time-locked to the presentation of food stimuli. Epochs were 1,300 ms long, including a 100 ms baseline period before stimulus onset, and a post-stimulus window of 1,300 ms. The data were manually screened for artefacts, and an ICA was again conducted to identify and remove components containing eye artefacts. The data were screened once more to eliminate any residual artefacts and signals exceeding ±200 μV.

#### 3.1.6 Decoding Analysis

The decoding parameters and overall approach were the same as those of Experiment 1 (see Section 2.1.6). For Experiment 2, however, three analyses were performed, all using the same EEG epochs from the decision stage – one predicting taste ratings, one predicting health ratings, and one predicting decision strength (ratings reflecting willingness to eat the foods presented in the decision stage). As in Experiment 1, an SVR approach was used for each analysis. The only difference was that health and taste rating scales in Experiment 2 were discrete (which reduces the variance, but otherwise does not matter for the SVR; Bode et al., 2014). The ratings for decision strength were again a continuous measure.

### 3.2 Results

Mean ratings for the full sample were 3.02 (*SE* = 0.09) for taste, 2.48 (*SE* = 0.12) for health, and -3.23 (*SD* = 6.55) for decision strength. A linear mixed effects model with participant as a random effect and the attributes health and taste as fixed effects found that decision strength was significantly predicted by health, β = 4.49, *t*(4567) = 7.55, *p* < .001, and taste, β = 43.99, *t*(4624) = 50.94, *p* < .001. For each participant, the correlation (Pearson’s *r*) between health and taste ratings ranged between -.27 and .56 (*mean r* = .13, *SD* = .19). Overall, there was a significant correlation between taste and health, *r*(87) = .28, *p* = .009.

The decoding of taste began to exceed chance from 740 ms after image onset, with a second cluster starting at 1120 ms (see Figure 3A). Of these, only the second cluster remained significant after correction for multiple comparisons; however, the large and sustained extent of the first cluster strongly suggests that this reflected meaningful information. For health, decoding performance was significantly above chance from 530 ms after image onset (see Figure 3B). Interestingly, even when the first cluster of taste decoding is considered meaningful, health decoding still appeared ∼ 200 ms earlier than taste decoding.

In the final analysis, the decoding of decision strength was possible from 830 ms after image onset. This result shows that fine-grained information about decision strength was indeed represented after taste and health representations, suggesting that it is at least possible that both sources of information contributed to computing the decision (see Figure 4). For test statistics from each analysis at each timepoint, see Supplementary Table S8.

**Figure 4.**
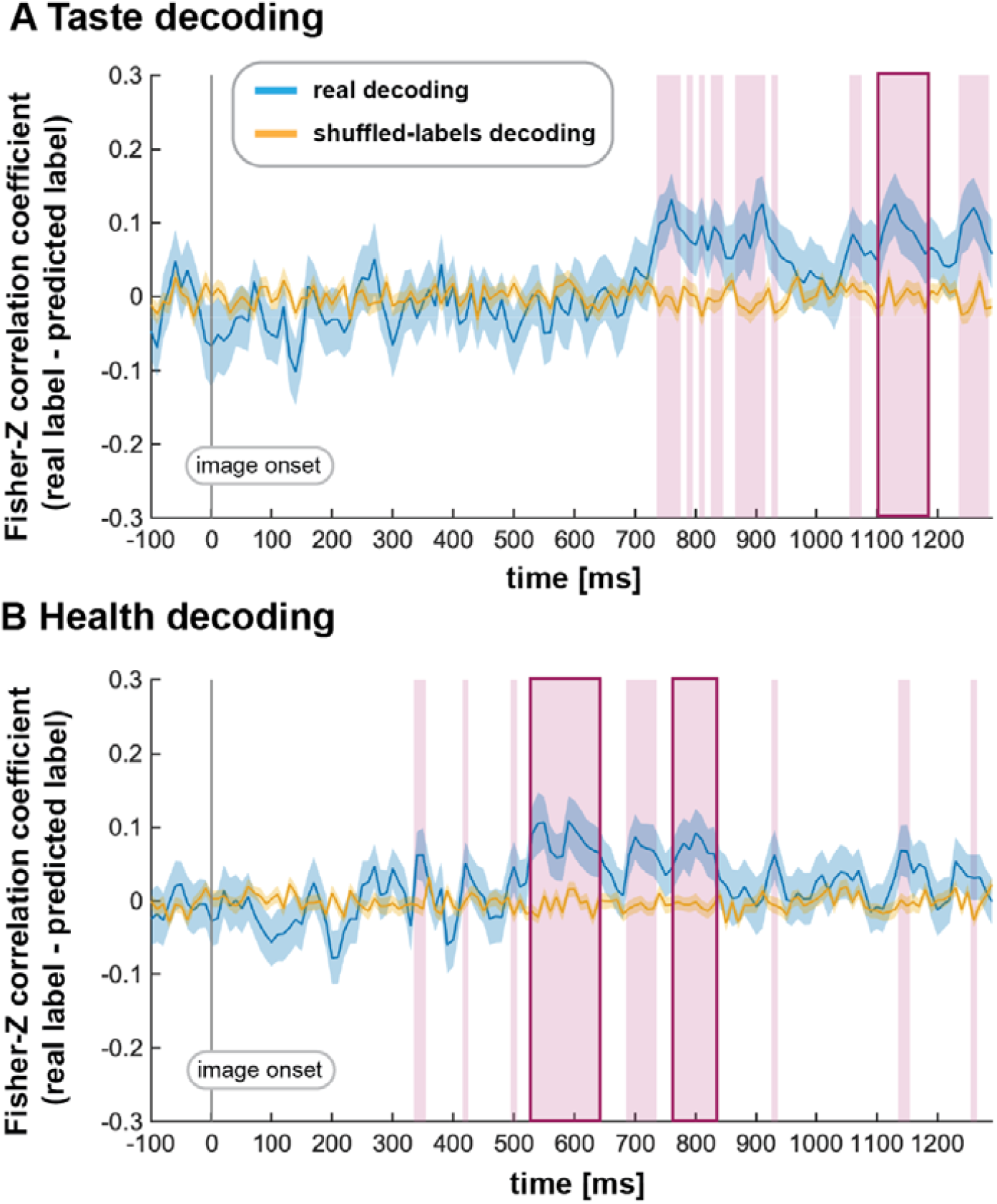
Decoding results from Experiment 2 for A) taste (*n* = 55) and B) health ratings (*n* = 85). Blue lines represent Fisher-transformed correlation coefficients between predicted and true labels (taste and health ratings), while yellow lines represent decoding results based on shuffled labels (empirical chance distribution). Pink vertical bars highlight time points at which the decoding results significantly differed from the shuffled decoding results (*p* < .05); bolded pink bars represent time windows that remained significant after corrections for multiple comparisons.

**Figure 5.**
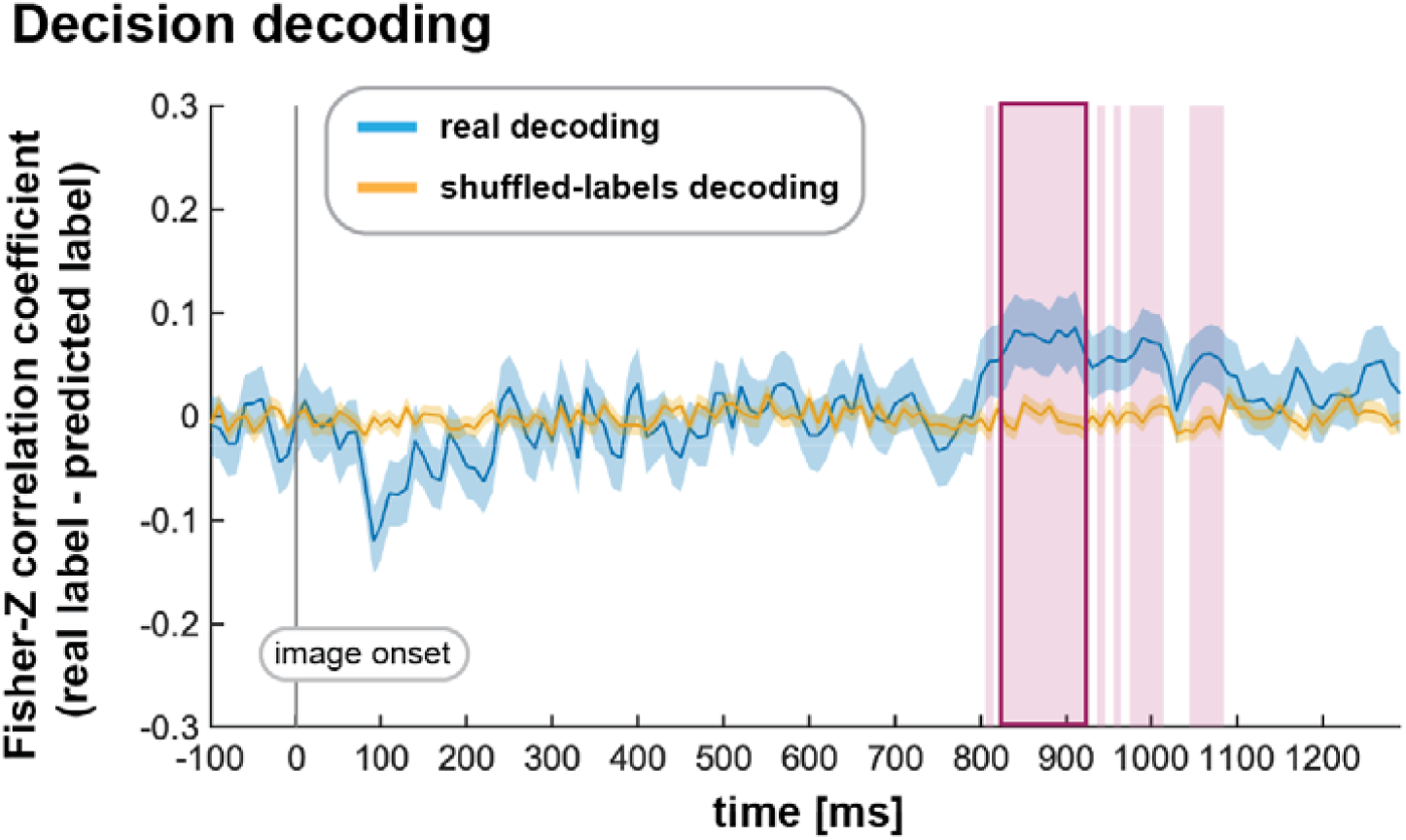
Decoding results from Experiment 2 for decision strength (*n* = 89). Blue lines represent Fisher-transformed correlation coefficients between predicted and true labels (taste and health ratings), while yellow lines represent decoding results based on shuffled labels (empirical chance distribution). Pink vertical bars highlight time points at which the decoding results significantly differed from the shuffled decoding results (*p* < .05); bolded pink bars represent time windows that remained significant after corrections for multiple comparisons.

## Discussion

We conducted two experiments to investigate the neural representations of taste and health attributes. In Experiment 1, we showed that patterns of neural activity in the first 1200 ms after a food image appeared predicted trial-wise ratings of these attributes when participants explicitly considered the tastiness and healthiness of various foods. In Experiment 2, we investigated whether representations of the same attributes could still be decoded from neural activity during a dietary decision, with no direct instruction to consider tastiness or healthiness. We found that it was possible to decode both health and taste ratings, as well as decision strength, from patterns of neural activity occurring in the first second after a food image appeared, even when participants were not instructed to explicitly consider these aspects.

The results from both experiments show that even subtle differences in the subjective perception of attributes of food stimuli are represented in EEG data, following on from previous ERP studies showing differences in average amplitudes between high- versus low-calorie and tasty versus non-tasty foods and drinks (Schwab et al., 2017; Schubert et al., 2021). For the first time, we also demonstrate that fine differences in health perception could be decoded from early neural activity. While there is evidence that dietary decisions are strongly driven by taste (Tepper & Trail, 1998; Pollard et al., 1998; Markovina et al., 2015; Morawetz et al., 2020), our results directly show that health information is also present in the brain at a similar time, before people make dietary decisions and when they are not explicitly instructed to think about health. Moreover, in our study, both taste and health ratings predicted food decisions, suggesting that both attributes are integrated for making the final choice. This further implies that interventions, such as health warning messages (Rosenblatt et al., 2018; Cohen & Lesser, 2016), targeted at modifying health attribute processing, could realistically change neural representations at early stages of dietary decisions.

In Experiment 1, in which participants explicitly processed either taste or health in separate trials, our results showed an earlier decoding onset for taste compared to health. This supports previous findings that taste is more salient (Motoki et al., 2018) and is processed more automatically (Liberman & Trope, 2008), and quickly than health (Sullivan et al., 2015). Interestingly, in Experiment 2, in which participants were not explicitly asked to consider tastiness or healthiness during dietary decision-making, health decoding onset was earlier than that of taste (in fact, health became decodable around the same time as taste in Experiment 1). This is surprising, considering the previous studies point to a general advantage for taste processing (e.g., Sullivan et al., 2015; Lim et al., 2018). One possible explanation for this reversal is that health processing was given priority because our food stimuli varied substantially in perceived healthiness. In contrast, most foods were rated as relatively tasty, and hence the variance of taste ratings was smaller than that of health ratings. In consequence, prioritising the quick assessment of health attributes of (mostly) tasty foods might have been a valuable strategy in this study. Future experiments could include more “not tasty” foods to test whether this would change the temporal order of decoding; however, in reality, a large number of “not tasty” options might prove difficult to find.

Another potential factor is that in Experiment 1, the food images were all of unpackaged foods, whereas many of the snack food images in Experiment 2 were presented in packaging (e.g., branded packets of chips). These packaged food stimuli may have been less likely to prompt the imagination of consumption (Muñoz-Vilches et al., 2020), leading to weaker representations of taste compared to the unpackaged stimuli in Experiment 1. Finally, another explanation for the lack of earlier taste decoding in Experiment 2 might be the lower statistical power relative to health decoding. However, given that the taste decoding analysis still comprised data from more than 50 participants, and clearly showed significant results later in the trial, this explanation is rather unlikely.

Regardless of the exact reason for why health attribute processing seemed to be temporally prioritised over taste processing, the findings of Experiment 2 strongly suggest that although correlated, both attributes were independently important for decision-making. Each attribute individually predicted the final dietary decisions in our regression analysis, and showed a different neural decoding time course. This suggests that the attributes were independent, at least to an extent. Finally, decoding of both attributes preceded the decoding of decision outcomes in time. While this is not direct evidence for causality, it suggests that these neural representations directly reflect the process by which participants inform their food decisions.

In conclusion, our results show that representations of taste and health attributes of foods can be decoded from brain activity within the first second of cognitive processing, both when participants are explicitly instructed to reflect on them, and when these attributes tacitly inform dietary decision-making. This constitutes a promising finding for public health interventions. Many recently suggested interventions, such as health warning labels at the point-of-sale, rely on their ability to quickly activate an implicit health goal, which can then be experienced as being in conflict with the perceived healthiness of the considered food item (Papies et al., 2016; Rosenblatt et al., 2018). For this mechanism to be effective, the healthiness of the food has to be perceived quickly, and crucially before the decision-process has concluded. Our results provide direct neural evidence that this is indeed the case (at least in situations in which a variety of items needs to be considered). Future research must now address whether having engaged with one’s own health perception beforehand (as in our Experiment 2) is a necessary condition, and whether interventions (such as nutrition labelling; Lim et al., 2018, or general and specific health warning messages; Rosenblatt et al., 2018a,b; Schubert et al., 2020) can alter the temporal processing dynamics – or directly the neuro-cognitive representation – of health and taste in the brain during dietary decision-making.

