## Supplementary Material for "Decoding explicit and implicit representations of health and taste attributes of foods in the human brain"

**Supplementary Material 1 - Experiment 1**

**Questionnaires**

Participants completed the Dutch Eating Behaviour Questionnaire (DEBQ; Van Strien et al., 1986), which consists of three scales measuring restrained eating (i.e., the tendency to restrict food intake), emotional eating (i.e., the tendency to eat in response to emotional changes), and external eating (i.e., the tendency to eat in response to external food cues). They also completed the Yale Food Addiction Scale (YFAS; Gearhardt et al., 2009), which measures clinically significant addiction to food.

**Results.** The sample had an average BMI of 23.75 (*SD* = 4.66; range 18.26 to 36.63), which falls within the normal range (Nuttall, 2015). Average self-reported hunger levels were 5.19 (scale from 1 “not hungry at all” to 9 “very hungry”; *SD* = 1.85), indicating that participants were slightly hungry. On average, participants reported that 8.92 hours (*SD* = 5.32) had passed since their last meal, and 7.27 hours (*SD* = 6.05) since their last snack. This suggests that most participants adhered to the condition of not eating for four hours before the experiment. The questionnaire showed that our sample significantly exceeded norms of all three scales of the DEBQ (Table S1). Note, however, that the questionnaire is normed for a German sample, and no Australian norms currently exist (Nagl et al., 2016). On the YFAS (Table S2), our sample did not appear to differ greatly from norms taken from North American undergraduates (Gearhardt et al., 2009), except on the “Tolerance” and “Continued use despite problems” scales. Three participants met the criteria for clinically significant food addiction but were still included in the sample.

**Supplementary Table S1.**

Sample means from Experiment 1 for the Dutch Eating Behavior (DEBQ) questionnaire, along with norms and statistical tests for comparison.

|  | M | SD | M_Norm_ | SD_Norm_ | *t* | *p* |
| --- | --- | --- | --- | --- | --- | --- |
| DEBQ-Restraint | 2.81 | 0.75 | 2.23 | 0.87 | 4.42 | < .001 |
| DEBQ-Emotional | 2.65 | 0.84 | 1.70 | 0.76 | 6.82 | < .001 |
| DEBQ-External | 3.19 | 0.58 | 2.57 | 0.74 | 7.04 | < .001 |

*Note:* Norms are taken from Nagl et al. (2016) for a German sample. Ratings were given on a scale from 1 to 5. The DEBQ measures restrained eating, emotional eating, and external eating.

**Supplementary Table S2.**

Results for the Yale Food Addiction Scale (YFAS) questionnaire, including sample percentages on each component, and norm percentages for comparison.

|  | Percentage | Norm Percentage |
| --- | --- | --- |
| Diagnosis of Food Dependence | 8.1% | 11.6% |
| Withdrawal | 13.5% | 16.3% |
| Tolerance | 89.2% | 13.5% |
| Continued Use Despite Problems | 73.0% | 28.3% |
| Important Activities Given Up | 24.3% | 10.3% |
| Large Amounts of Time Spent | 10.81% | 24.0% |
| Loss of Control | 13.51% | 21.7% |
| Have Tried Unsuccessfully to Cut Down or Worried About Cutting Down | 86.5% | 71.3% |
| Clinically Significant Impairment | 8.1% | 14.0% |

**Supplementary Table S3.**

*Test statistics for each taste and health multivariate pattern analysis in Experiment 1, including p- and t-values, and Cohen’s d. Results have not been corrected for multiple comparisons.*

|  | Taste | | | Health | | |
| --- | --- | --- | --- | --- | --- | --- |
| Time Step | *t* | *p* | *d* | *t* | *p* | *d* |
| 1 | 2.21 | 0.02** | 4.03 | -0.10 | 0.54* | -0.11 |
| 2 | 0.62 | 0.27 | 1.24 | -0.43 | 0.66 | -0.49 |
| 3 | -2.52 | 0.99 | -4.74 | -0.02 | 0.51 | -0.03 |
| 4 | -0.96 | 0.83 | -1.81 | 0.11 | 0.46 | 0.17 |
| 5 | 0.50 | 0.31 | 1.04 | -0.45 | 0.67 | -0.64 |
| 6 | 0.66 | 0.26 | 1.10 | -0.16 | 0.56 | -0.22 |
| 7 | -0.43 | 0.67 | -0.71 | 0.75 | 0.23 | 1.06 |
| 8 | -0.35 | 0.64 | -0.66 | -0.07 | 0.53 | -0.08 |
| 9 | -0.13 | 0.55 | -0.22 | -0.51 | 0.69 | -0.64 |
| 10 | 0.93 | 0.18 | 1.44 | -0.76 | 0.77 | -1.06 |
| 11 | 0.49 | 0.31 | 0.70 | -0.71 | 0.76 | -0.97 |
| 12 | 0.93 | 0.18 | 1.43 | -1.50 | 0.93 | -1.45 |
| 13 | 1.36 | 0.09* | 2.13 | -1.24 | 0.89 | -1.58 |
| 14 | 0.89 | 0.19 | 1.53 | -0.93 | 0.82 | -1.64 |
| 15 | 0.24 | 0.40 | 0.41 | -0.31 | 0.62 | -0.45 |
| 16 | -0.31 | 0.62 | -0.53 | 0.60 | 0.28 | 0.77 |
| 17 | -0.52 | 0.70 | -0.89 | 0.87 | 0.19 | 1.02 |
| 18 | -1.34 | 0.91 | -2.18 | -0.05 | 0.52 | -0.06 |
| 19 | -0.75 | 0.77 | -0.98 | 0.18 | 0.43 | 0.24 |
| 20 | -1.77 | 0.96 | -2.85 | 1.23 | 0.11 | 1.51 |
| 21 | -1.69 | 0.95 | -2.87 | 1.35 | 0.09* | 1.48 |
| 22 | -0.10 | 0.54 | -0.18 | 1.31 | 0.10 | 1.21 |
| 23 | 0.94 | 0.18 | 1.52 | -0.38 | 0.65 | -0.64 |
| 24 | 0.97 | 0.17 | 1.43 | -0.62 | 0.73 | -0.77 |
| 25 | 2.21 | 0.02** | 2.90 | 0.36 | 0.36 | 0.49 |
| 26 | 0.98 | 0.17 | 1.36 | -0.71 | 0.76 | -1.39 |
| 27 | 0.48 | 0.32 | 0.67 | -0.88 | 0.81 | -1.24 |
| 28 | -0.02 | 0.51 | -0.03 | -1.45 | 0.92 | -2.07 |
| 29 | 0.60 | 0.27 | 1.19 | -1.10 | 0.86 | -1.83 |
| 30 | 0.45 | 0.33 | 0.87 | -0.78 | 0.78 | -1.06 |
| 31 | 0.84 | 0.20 | 1.71 | -0.84 | 0.80 | -1.05 |
| 32 | 1.19 | 0.12 | 2.23 | -1.01 | 0.84 | -1.15 |
| 33 | 0.92 | 0.18 | 1.62 | -0.12 | 0.55 | -0.12 |
| 34 | 0.39 | 0.35 | 0.65 | 0.05 | 0.48 | 0.06 |
| 35 | 0.47 | 0.32 | 0.91 | -0.91 | 0.81 | -1.13 |
| 36 | 1.13 | 0.13 | 1.67 | -0.97 | 0.83 | -1.09 |
| 37 | 1.48 | 0.07* | 2.69 | -1.01 | 0.84 | -1.40 |
| 38 | 0.41 | 0.34 | 0.77 | -0.18 | 0.57 | -0.20 |
| 39 | 1.15 | 0.13 | 2.22 | 0.47 | 0.32 | 0.47 |
| 40 | 1.36 | 0.09* | 2.24 | 0.01 | 0.50 | 0.01 |
| 41 | 2.04 | 0.02** | 3.42 | -0.24 | 0.60 | -0.31 |
| 42 | 2.51 | 0.01** | 4.28 | 0.42 | 0.34 | 0.52 |
| 43 | 1.90 | 0.03** | 2.74 | 0.48 | 0.32 | 0.62 |
| 44 | 1.26 | 0.11 | 1.92 | 0.41 | 0.34 | 0.56 |
| 45 | 0.47 | 0.32 | 0.74 | 0.14 | 0.44 | 0.17 |
| 46 | 1.06 | 0.15 | 1.96 | 0.12 | 0.45 | 0.13 |
| 47 | -0.08 | 0.53 | -0.19 | -0.57 | 0.71 | -0.73 |
| 48 | 0.86 | 0.20 | 1.87 | -0.04 | 0.51 | -0.05 |
| 49 | 1.35 | 0.09* | 2.48 | 0.50 | 0.31 | 0.52 |
| 50 | 2.01 | 0.03** | 3.27 | 0.81 | 0.21 | 0.96 |
| 51 | 1.31 | 0.10 | 2.32 | 0.48 | 0.32 | 0.43 |
| 52 | 0.97 | 0.17 | 1.79 | 1.28 | 0.10 | 1.53 |
| 53 | 1.02 | 0.16 | 1.95 | 0.80 | 0.22 | 0.87 |
| 54 | 0.70 | 0.24 | 1.47 | 0.53 | 0.30 | 0.68 |
| 55 | 0.20 | 0.42 | 0.43 | 0.71 | 0.24 | 0.94 |
| 56 | 0.64 | 0.26 | 1.21 | 1.35 | 0.09* | 2.87 |
| 57 | 0.14 | 0.44 | 0.26 | 1.49 | 0.07* | 2.70 |
| 58 | 0.40 | 0.34 | 0.80 | 1.34 | 0.09* | 2.17 |
| 59 | 0.58 | 0.28 | 0.92 | 0.78 | 0.22 | 0.88 |
| 60 | -0.10 | 0.54 | -0.20 | 0.43 | 0.34 | 0.56 |
| 61 | 0.45 | 0.33 | 0.85 | 0.35 | 0.36 | 0.40 |
| 62 | 0.26 | 0.40 | 0.48 | 0.50 | 0.31 | 0.57 |
| 63 | 1.53 | 0.07* | 3.01 | 0.65 | 0.26 | 0.76 |
| 64 | 1.04 | 0.15 | 1.66 | 2.35 | 0.01** | 2.97 |
| 65 | 1.04 | 0.15 | 1.83 | 3.19 | 0.00*** | 3.54 |
| 66 | 0.99 | 0.17 | 2.05 | 4.27 | 0.00*** | 4.46 |
| 67 | 0.91 | 0.18 | 1.92 | 4.03 | 0.00*** | 3.87 |
| 68 | 1.45 | 0.08* | 2.70 | 3.81 | 0.00*** | 4.55 |
| 69 | 1.58 | 0.06* | 2.84 | 3.11 | 0.00*** | 3.40 |
| 70 | 1.23 | 0.11 | 2.47 | 2.70 | 0.01** | 3.64 |
| 71 | 0.77 | 0.22 | 1.48 | 1.97 | 0.03** | 3.36 |
| 72 | 0.45 | 0.33 | 0.86 | 1.94 | 0.03** | 2.37 |
| 73 | 0.74 | 0.23 | 1.47 | 1.83 | 0.04** | 2.83 |
| 74 | 1.61 | 0.06* | 2.49 | 2.09 | 0.02** | 3.27 |
| 75 | 2.66 | 0.01** | 4.34 | 2.14 | 0.02** | 2.86 |
| 76 | 2.38 | 0.01** | 4.13 | 1.67 | 0.05* | 3.08 |
| 77 | 3.02 | 0.00*** | 4.59 | 1.53 | 0.07* | 2.01 |
| 78 | 2.29 | 0.01** | 3.40 | 1.62 | 0.06* | 2.75 |
| 79 | 1.95 | 0.03** | 2.66 | 2.34 | 0.01** | 2.63 |
| 80 | 3.56 | 0.00*** | 5.07 | 2.20 | 0.02** | 2.77 |
| 81 | 2.55 | 0.01** | 4.37 | 3.39 | 0.00*** | 4.44 |
| 82 | 2.18 | 0.02** | 3.81 | 2.98 | 0.00*** | 4.09 |
| 83 | 1.42 | 0.08* | 2.57 | 3.04 | 0.00*** | 4.35 |
| 84 | 1.53 | 0.07* | 2.26 | 3.44 | 0.00*** | 3.71 |
| 85 | 1.51 | 0.07* | 2.10 | 3.50 | 0.00*** | 4.05 |
| 86 | 3.10 | 0.00*** | 3.99 | 2.49 | 0.01** | 2.85 |
| 87 | 2.81 | 0.00*** | 3.67 | 2.06 | 0.02** | 2.78 |
| 88 | 3.59 | 0.00*** | 5.31 | 2.16 | 0.02** | 2.11 |
| 89 | 2.48 | 0.01** | 3.69 | 1.50 | 0.07* | 2.44 |
| 90 | 2.58 | 0.01** | 3.94 | 1.32 | 0.10 | 1.75 |
| 91 | 2.14 | 0.02** | 2.85 | 1.77 | 0.04** | 2.48 |
| 92 | 1.50 | 0.07* | 2.14 | 1.34 | 0.09* | 1.91 |
| 93 | 1.62 | 0.06* | 2.24 | 2.22 | 0.02** | 2.36 |
| 94 | 1.70 | 0.05* | 2.16 | 1.86 | 0.04** | 2.09 |
| 95 | 1.52 | 0.07* | 1.73 | 1.99 | 0.03** | 2.13 |
| 96 | 2.31 | 0.01** | 2.85 | 2.97 | 0.00*** | 3.77 |
| 97 | 1.34 | 0.09* | 1.96 | 2.83 | 0.00*** | 3.33 |
| 98 | 1.91 | 0.03** | 2.59 | 2.71 | 0.01*** | 3.28 |
| 99 | 1.03 | 0.15 | 1.29 | 3.61 | 0.00*** | 4.16 |
| 100 | 1.36 | 0.09* | 2.28 | 3.98 | 0.00*** | 4.27 |
| 101 | 1.24 | 0.11 | 2.18 | 3.11 | 0.00*** | 3.47 |
| 102 | 1.78 | 0.04** | 2.57 | 2.36 | 0.01** | 2.35 |
| 103 | 0.95 | 0.18 | 1.26 | 2.53 | 0.01** | 2.49 |
| 104 | 1.52 | 0.07* | 2.55 | 3.39 | 0.00*** | 3.12 |
| 105 | 1.55 | 0.07* | 2.26 | 1.75 | 0.04** | 1.89 |
| 106 | 1.67 | 0.05* | 2.46 | 2.60 | 0.01** | 2.59 |
| 107 | 0.81 | 0.21 | 1.30 | 2.27 | 0.01** | 2.59 |
| 108 | 1.07 | 0.14 | 1.84 | 2.45 | 0.01** | 3.28 |
| 109 | 1.09 | 0.14 | 1.76 | 1.69 | 0.05* | 2.46 |
| 110 | 1.12 | 0.13 | 2.33 | 0.95 | 0.17 | 1.28 |
| 111 | 1.43 | 0.08* | 3.04 | 1.20 | 0.12 | 1.76 |
| 112 | 1.09 | 0.14 | 1.99 | 1.67 | 0.05* | 2.20 |
| 113 | 0.97 | 0.17 | 1.76 | 0.49 | 0.31 | 0.53 |
| 114 | 0.67 | 0.25 | 1.20 | 1.02 | 0.16 | 1.19 |
| 115 | 0.40 | 0.34 | 0.80 | 0.59 | 0.28 | 0.70 |
| 116 | 0.16 | 0.44 | 0.31 | 1.35 | 0.09* | 1.57 |
| 117 | 0.22 | 0.41 | 0.44 | 1.55 | 0.07* | 1.95 |
| 118 | 1.14 | 0.13 | 2.40 | 0.65 | 0.26 | 0.76 |
| 119 | 1.60 | 0.06* | 2.91 | 0.64 | 0.26 | 0.71 |
| 120 | 1.30 | 0.10 | 2.29 | 1.01 | 0.16 | 1.25 |
| 121 | 0.74 | 0.23 | 1.55 | 1.00 | 0.16 | 1.49 |
| 122 | 0.28 | 0.39 | 0.66 | 1.27 | 0.11 | 1.79 |
| 123 | 0.66 | 0.26 | 1.48 | 0.54 | 0.30 | 0.75 |
| 124 | 1.40 | 0.08* | 3.14 | 1.03 | 0.16 | 1.40 |
| 125 | 1.49 | 0.07* | 3.02 | 0.41 | 0.34 | 0.44 |
| 126 | 1.48 | 0.07* | 2.72 | 0.59 | 0.28 | 0.67 |
| 127 | 1.16 | 0.13 | 2.02 | 0.89 | 0.19 | 1.15 |
| 128 | 1.05 | 0.15 | 1.98 | -0.10 | 0.54 | -0.12 |
| 129 | 0.87 | 0.20 | 1.98 | 0.09 | 0.46 | 0.12 |
| 130 | 0.69 | 0.25 | 1.28 | 0.41 | 0.34 | 0.62 |
| 131 | 1.14 | 0.13 | 2.29 | 0.50 | 0.31 | 0.64 |

Note: * *p* < .10, ** *p* < .05, *** *p* < .001

**Supplementary Material 2 – Experiment 2**

Participants completed the DEBQ (Van Strien et al., 1986; see Section 2.1.3) and the Food Choice Questionnaire (FCQ; Pollard et al., 1998), which measures the importance of nine factors underlying an individual’s food choice, such as convenience, health, and ethical concern, on a four-point scale (1 = not at all important; 4 = very important).

**Results**

The average BMI of the sample was 21.02 (*SD* = 2.24; range 16.65 to 27.77; note that 12 participants chose not to report their height/weight and are therefore excluded from this average), which falls within the “normal” range (Nuttall, 2015). On average, participants had fasted for 6.95 hours (*SD* = 4.66) and gave hunger ratings of 4.63 (scale from 1 = not at all hungry to 8 = extremely hungry; *SD* = 1.70). As in Experiment 1, the sample significantly exceeded the norms (Nagl et al., 2016) on all three dimensions of the DEBQ (see Table S4). As mentioned above, the norms from Nagl et al. (2016) are taken from a German sample – as the Australian samples in Experiments 1 and 2 both exceeded the norms, this can likely be attributed to cultural differences. On the FCQ, our sample significantly exceeded the norms (Pollard et al., 1998) for mood, convenience, price, familiarity, and ethical concern (see Table S5).

**Supplementary Table S4.**

*Sample means from Experiment 2 for the Dutch Eating Behavior (DEBQ) questionnaire, along with norms and statistical tests for comparison.*

|  | M | SD | M_Norm_ | SD_Norm_ | *t* | *p* |
| --- | --- | --- | --- | --- | --- | --- |
| DEBQ-Restraint | 2.77 | 1.35 | 2.23 | 0.87 | 3.78 | < .001 |
| DEBQ-Emotional | 2.60 | 1.18 | 1.70 | 0.76 | 7.25 | < .001 |
| DEBQ-External | 3.17 | 0.87 | 2.57 | 0.74 | 6.55 | < .001 |

*Note:* Norms are taken from Nagl et al. (2016) for a German sample. Ratings were given on a scale from 1 to 5. The DEBQ measures restrained eating, emotional eating, and external eating.

**Supplementary Table S5.**

*Sample means from Experiment 2 for the Food Choice Questionnaire (FCQ), along with norms and statistical tests for comparison.*

|  | M | SD | M_Norm_ | SD_Norm_ | *t* | *p* |
| --- | --- | --- | --- | --- | --- | --- |
| Health | 2.95 | 0.78 | 2.82 | 0.72 | 1.60 | .114 |
| Mood | 2.67 | 0.80 | 2.10 | 0.73 | 6.73 | < .001 |
| Convenience | 2.99 | 0.67 | 2.75 | 0.80 | 3.29 | .001 |
| Sensory Appeal | 2.88 | 0.55 | 2.99 | 0.63 | -1.80 | .075 |
| Natural Content | 2.81 | 0.69 | 2.45 | 0.86 | -0.01 | .999 |
| Price | 2.97 | 0.69 | 2.83 | 0.80 | 2.03 | .045 |
| Weight Control | 2.48 | 0.90 | 2.38 | 0.87 | 0.95 | .345 |
| Familiarity | 2.41 | 0.73 | 1.76 | 0.68 | 8.43 | < .001 |
| Ethical Concern | 2.07 | 0.82 | 1.86 | 0.78 | 2.34 | .021 |

*Note:* Norms are taken from Pollard et al. (1998) for a sample from the UK. Ratings were given on a scale from 1 to 4.

**Supplementary Table S6.**

*Test statistics for the taste, health, and decision strength multivariate pattern analysis results in Experiment 2, including p- and t-values, and Cohen’s d. Results have not been corrected for multiple comparisons.*

|  | Taste | | | Health | | | Decision Strength | | |
| --- | --- | --- | --- | --- | --- | --- | --- | --- | --- |
| Time Step | *t* | *p* | *d* | *t* | *p* | *d* | *t* | *p* | *d* |
| 1 | -0.87 | 0.805 | -1.09 | -0.61 | 0.727 | -0.90 | -0.07 | 0.528 | -0.09 |
| 2 | -1.05 | 0.852 | -1.40 | -0.15 | 0.560 | -0.25 | -0.72 | 0.764 | -0.92 |
| 3 | -0.13 | 0.552 | -0.20 | -0.58 | 0.717 | -0.87 | -0.33 | 0.628 | -0.45 |
| 4 | 0.44 | 0.329 | 0.65 | 0.72 | 0.237 | 1.02 | -0.75 | 0.771 | -0.91 |
| 5 | 0.52 | 0.304 | 0.68 | 0.74 | 0.230 | 0.97 | 0.37 | 0.357 | 0.46 |
| 6 | 0.16 | 0.437 | 0.22 | 0.93 | 0.177 | 1.18 | 0.88 | 0.190 | 1.14 |
| 7 | 0.88 | 0.191 | 1.13 | 0.61 | 0.271 | 0.87 | 0.71 | 0.240 | 0.94 |
| 8 | 0.74 | 0.232 | 1.14 | -0.22 | 0.585 | -0.31 | -0.73 | 0.767 | -0.90 |
| 9 | -0.34 | 0.633 | -0.43 | -0.21 | 0.585 | -0.29 | -1.62 | 0.945 | -2.03 |
| 10 | -1.30 | 0.900 | -2.33 | -0.65 | 0.740 | -0.83 | -0.82 | 0.794 | -1.02 |
| 11 | -1.31 | 0.902 | -2.01 | -0.83 | 0.796 | -1.17 | -0.13 | 0.552 | -0.20 |
| 12 | -1.16 | 0.875 | -1.87 | -0.66 | 0.744 | -1.00 | 0.22 | 0.414 | 0.35 |
| 13 | -1.09 | 0.860 | -1.84 | 0.19 | 0.427 | 0.26 | 0.16 | 0.436 | 0.25 |
| 14 | -0.73 | 0.766 | -0.87 | -0.09 | 0.537 | -0.14 | -0.12 | 0.547 | -0.18 |
| 15 | -0.43 | 0.667 | -0.58 | 0.20 | 0.421 | 0.29 | 0.20 | 0.421 | 0.26 |
| 16 | -0.54 | 0.705 | -0.80 | -0.29 | 0.614 | -0.41 | -1.00 | 0.840 | -1.48 |
| 17 | -0.56 | 0.712 | -0.76 | -0.40 | 0.656 | -0.54 | -0.49 | 0.686 | -0.62 |
| 18 | 1.15 | 0.128 | 1.23 | -0.84 | 0.800 | -1.09 | -0.22 | 0.587 | -0.28 |
| 19 | -0.56 | 0.710 | -0.74 | -1.16 | 0.876 | -1.61 | -1.50 | 0.931 | -1.81 |
| 20 | -1.36 | 0.910 | -1.81 | -1.29 | 0.899 | -1.72 | -3.72 | 1.000 | -4.72 |
| 21 | -1.19 | 0.881 | -1.33 | -1.72 | 0.955 | -2.31 | -2.86 | 0.997 | -3.87 |
| 22 | -1.33 | 0.905 | -1.61 | -1.48 | 0.928 | -2.17 | -2.19 | 0.984 | -2.86 |
| 23 | -0.79 | 0.783 | -1.06 | -0.92 | 0.821 | -1.40 | -1.95 | 0.973 | -2.56 |
| 24 | -1.84 | 0.964 | -2.61 | -1.62 | 0.945 | -2.33 | -2.23 | 0.986 | -3.14 |
| 25 | -1.81 | 0.962 | -2.46 | -0.94 | 0.825 | -1.36 | -0.45 | 0.674 | -0.61 |
| 26 | -1.28 | 0.897 | -1.84 | -0.56 | 0.710 | -0.83 | -1.34 | 0.908 | -1.68 |
| 27 | 0.38 | 0.352 | 0.50 | 0.21 | 0.415 | 0.29 | -1.93 | 0.971 | -2.39 |
| 28 | -0.22 | 0.587 | -0.28 | 0.74 | 0.232 | 1.07 | -1.95 | 0.973 | -2.47 |
| 29 | -0.12 | 0.546 | -0.16 | 0.04 | 0.485 | 0.05 | -0.15 | 0.560 | -0.18 |
| 30 | -0.93 | 0.822 | -1.36 | -0.28 | 0.610 | -0.42 | -0.56 | 0.711 | -0.69 |
| 31 | -0.49 | 0.686 | -0.64 | -2.33 | 0.989 | -3.47 | -1.21 | 0.886 | -1.57 |
| 32 | -0.75 | 0.771 | -1.09 | -1.86 | 0.967 | -2.79 | -1.10 | 0.863 | -1.54 |
| 33 | -1.76 | 0.958 | -2.05 | -0.53 | 0.703 | -0.78 | -1.79 | 0.962 | -2.51 |
| 34 | -0.05 | 0.518 | -0.06 | -1.08 | 0.858 | -1.49 | -1.51 | 0.932 | -1.99 |
| 35 | 0.11 | 0.457 | 0.14 | -0.66 | 0.744 | -0.94 | 0.64 | 0.261 | 0.92 |
| 36 | 0.38 | 0.353 | 0.54 | 0.87 | 0.195 | 1.14 | 0.85 | 0.199 | 1.08 |
| 37 | 0.73 | 0.234 | 1.07 | 1.30 | 0.098* | 1.52 | 0.36 | 0.361 | 0.49 |
| 38 | 1.21 | 0.116 | 1.89 | 1.45 | 0.076* | 1.73 | -0.26 | 0.601 | -0.36 |
| 39 | 0.13 | 0.449 | 0.20 | 0.44 | 0.330 | 0.60 | -0.96 | 0.830 | -1.28 |
| 40 | -1.08 | 0.858 | -1.54 | 0.56 | 0.288 | 0.71 | 0.09 | 0.464 | 0.11 |
| 41 | -1.11 | 0.865 | -1.68 | 1.56 | 0.062* | 1.83 | -0.16 | 0.564 | -0.21 |
| 42 | -0.73 | 0.764 | -1.04 | 1.09 | 0.139 | 1.41 | 0.29 | 0.386 | 0.44 |
| 43 | 0.00 | 0.500 | 0.00 | 0.50 | 0.308 | 0.66 | 0.18 | 0.428 | 0.25 |
| 44 | -0.22 | 0.588 | -0.30 | -0.94 | 0.826 | -1.23 | -1.38 | 0.915 | -1.88 |
| 45 | -0.46 | 0.677 | -0.64 | 1.87 | 0.032** | 2.61 | 1.15 | 0.126 | 1.38 |
| 46 | -0.11 | 0.545 | -0.15 | 1.89 | 0.031** | 2.82 | 0.23 | 0.409 | 0.32 |
| 47 | 0.22 | 0.413 | 0.30 | -0.21 | 0.582 | -0.30 | -0.58 | 0.718 | -0.77 |
| 48 | -0.65 | 0.740 | -0.95 | 0.07 | 0.470 | 0.10 | -1.03 | 0.848 | -1.41 |
| 49 | 1.22 | 0.114 | 1.72 | 0.85 | 0.198 | 1.34 | -0.86 | 0.805 | -1.20 |
| 50 | -0.42 | 0.663 | -0.66 | -1.90 | 0.970 | -2.91 | 0.74 | 0.230 | 0.97 |
| 51 | -0.08 | 0.532 | -0.12 | -1.15 | 0.873 | -1.80 | 1.13 | 0.130 | 1.63 |
| 52 | -0.56 | 0.711 | -0.77 | 0.18 | 0.429 | 0.24 | -0.18 | 0.573 | -0.26 |
| 53 | -0.75 | 0.771 | -0.87 | 1.67 | 0.049** | 1.99 | -0.30 | 0.617 | -0.37 |
| 54 | -0.31 | 0.620 | -0.38 | 0.54 | 0.295 | 0.72 | -0.53 | 0.701 | -0.66 |
| 55 | 0.11 | 0.455 | 0.16 | 0.26 | 0.396 | 0.37 | -0.97 | 0.834 | -1.49 |
| 56 | -0.36 | 0.642 | -0.56 | 0.47 | 0.318 | 0.76 | -1.46 | 0.926 | -2.01 |
| 57 | -0.38 | 0.646 | -0.53 | -0.65 | 0.741 | -0.97 | -0.06 | 0.525 | -0.07 |
| 58 | -0.80 | 0.786 | -1.13 | -0.61 | 0.727 | -0.95 | -1.11 | 0.865 | -1.29 |
| 59 | -0.32 | 0.625 | -0.40 | -0.42 | 0.663 | -0.65 | -0.79 | 0.785 | -1.04 |
| 60 | -0.96 | 0.828 | -1.39 | 0.46 | 0.324 | 0.63 | 0.76 | 0.226 | 0.98 |
| 61 | -1.74 | 0.956 | -2.66 | 1.86 | 0.033** | 2.54 | 0.35 | 0.362 | 0.45 |
| 62 | -0.87 | 0.806 | -1.30 | 0.45 | 0.326 | 0.68 | -0.85 | 0.800 | -1.01 |
| 63 | -0.13 | 0.553 | -0.20 | 1.10 | 0.138 | 1.62 | 0.78 | 0.218 | 1.01 |
| 64 | 0.11 | 0.456 | 0.17 | 2.85 | 0.003** | 4.10 | 0.20 | 0.421 | 0.29 |
| 65 | -0.16 | 0.565 | -0.26 | 2.93 | 0.002** | 4.93 | 0.11 | 0.454 | 0.16 |
| 66 | -1.17 | 0.877 | -1.74 | 2.73 | 0.004** | 3.85 | -0.48 | 0.682 | -0.55 |
| 67 | -0.84 | 0.797 | -1.18 | 2.10 | 0.020** | 3.18 | 0.76 | 0.226 | 0.95 |
| 68 | -0.27 | 0.605 | -0.43 | 1.82 | 0.036** | 2.08 | 1.03 | 0.153 | 1.39 |
| 69 | -0.64 | 0.737 | -1.22 | 2.07 | 0.021** | 2.92 | 0.70 | 0.242 | 0.94 |
| 70 | -1.08 | 0.857 | -1.78 | 2.47 | 0.008** | 3.59 | 0.05 | 0.482 | 0.06 |
| 71 | -0.03 | 0.511 | -0.05 | 1.99 | 0.025** | 3.17 | -1.06 | 0.853 | -1.46 |
| 72 | -0.01 | 0.503 | -0.01 | 2.34 | 0.011** | 3.46 | -0.29 | 0.615 | -0.46 |
| 73 | -0.37 | 0.642 | -0.71 | 2.02 | 0.023** | 2.90 | -0.71 | 0.760 | -0.95 |
| 74 | -0.05 | 0.518 | -0.10 | 2.54 | 0.006** | 3.57 | 0.82 | 0.207 | 1.15 |
| 75 | 0.53 | 0.297 | 1.04 | 1.88 | 0.032** | 2.51 | 0.31 | 0.379 | 0.47 |
| 76 | 0.39 | 0.351 | 0.73 | 1.28 | 0.102 | 1.82 | -0.06 | 0.525 | -0.09 |
| 77 | -0.14 | 0.557 | -0.28 | 0.98 | 0.165 | 1.44 | 0.95 | 0.172 | 1.20 |
| 78 | -0.44 | 0.668 | -1.03 | 0.42 | 0.339 | 0.51 | 0.68 | 0.249 | 0.91 |
| 79 | -0.77 | 0.778 | -1.58 | 0.77 | 0.222 | 0.99 | -0.49 | 0.686 | -0.61 |
| 80 | 0.15 | 0.440 | 0.30 | 2.18 | 0.016** | 2.80 | -0.01 | 0.502 | -0.01 |
| 81 | 0.54 | 0.296 | 1.08 | 2.89 | 0.002** | 3.58 | 0.68 | 0.249 | 0.84 |
| 82 | 0.60 | 0.277 | 1.23 | 2.37 | 0.010** | 3.00 | 0.28 | 0.391 | 0.39 |
| 83 | 0.07 | 0.472 | 0.16 | 2.35 | 0.010** | 3.17 | 0.66 | 0.257 | 0.95 |
| 84 | 1.14 | 0.130 | 2.40 | 2.15 | 0.017** | 3.05 | -0.05 | 0.519 | -0.07 |
| 85 | 2.68 | 0.005** | 5.03 | 1.18 | 0.121 | 1.64 | -0.21 | 0.582 | -0.30 |
| 86 | 2.76 | 0.004** | 4.58 | 1.32 | 0.095* | 1.59 | -0.71 | 0.761 | -1.05 |
| 87 | 3.68 | 0.000*** | 5.93 | 1.44 | 0.076* | 1.85 | -0.54 | 0.706 | -0.81 |
| 88 | 3.25 | 0.001** | 5.31 | 1.71 | 0.045** | 2.53 | -0.12 | 0.549 | -0.19 |
| 89 | 1.64 | 0.053* | 3.27 | 2.21 | 0.015** | 3.22 | 0.18 | 0.429 | 0.26 |
| 90 | 1.78 | 0.040** | 3.29 | 1.86 | 0.033** | 2.87 | 0.19 | 0.424 | 0.29 |
| 91 | 1.50 | 0.070* | 3.10 | 2.65 | 0.005** | 3.63 | 1.20 | 0.117 | 1.69 |
| 92 | 2.85 | 0.003** | 5.47 | 3.00 | 0.002** | 3.59 | 1.97 | 0.026** | 2.70 |
| 93 | 1.48 | 0.072* | 2.85 | 2.29 | 0.012** | 3.05 | 1.17 | 0.123 | 1.82 |
| 94 | 2.58 | 0.006** | 4.50 | 1.85 | 0.034** | 2.66 | 1.79 | 0.038** | 3.00 |
| 95 | 2.06 | 0.022** | 3.89 | 0.51 | 0.304 | 0.80 | 2.72 | 0.004** | 3.88 |
| 96 | 0.92 | 0.181 | 1.80 | 1.22 | 0.112 | 2.01 | 1.92 | 0.029** | 2.63 |
| 97 | 0.63 | 0.265 | 1.31 | 0.13 | 0.448 | 0.21 | 2.11 | 0.019** | 2.92 |
| 98 | 1.69 | 0.048** | 3.66 | 0.87 | 0.192 | 1.31 | 1.94 | 0.028** | 2.93 |
| 99 | 2.02 | 0.024** | 4.29 | 0.67 | 0.253 | 1.00 | 1.88 | 0.032** | 2.41 |
| 100 | 1.84 | 0.036** | 3.76 | 0.28 | 0.389 | 0.43 | 2.82 | 0.003** | 3.54 |
| 101 | 2.49 | 0.008** | 5.22 | -0.09 | 0.537 | -0.16 | 2.21 | 0.015** | 3.37 |
| 102 | 3.20 | 0.001** | 5.65 | -0.29 | 0.614 | -0.46 | 2.54 | 0.006** | 3.74 |
| 103 | 1.63 | 0.055* | 3.06 | 1.17 | 0.122 | 1.60 | 2.03 | 0.023** | 2.98 |
| 104 | 1.95 | 0.028** | 4.04 | 1.88 | 0.032** | 2.64 | 1.40 | 0.083* | 1.88 |
| 105 | 1.53 | 0.066* | 3.10 | 1.34 | 0.092* | 1.70 | 1.87 | 0.032** | 2.48 |
| 106 | 1.09 | 0.141 | 2.39 | 0.56 | 0.288 | 0.77 | 1.61 | 0.055* | 2.09 |
| 107 | 0.71 | 0.239 | 1.31 | -0.06 | 0.525 | -0.09 | 1.97 | 0.026** | 2.49 |
| 108 | -0.03 | 0.512 | -0.05 | 0.16 | 0.438 | 0.23 | 1.60 | 0.056* | 1.97 |
| 109 | 0.58 | 0.284 | 1.13 | 0.23 | 0.408 | 0.34 | 1.85 | 0.034** | 2.24 |
| 110 | 0.77 | 0.222 | 1.69 | 0.14 | 0.446 | 0.20 | 2.70 | 0.004** | 3.28 |
| 111 | 0.57 | 0.287 | 1.11 | 0.74 | 0.230 | 0.96 | 2.27 | 0.013** | 2.59 |
| 112 | 0.04 | 0.485 | 0.08 | 0.71 | 0.240 | 1.03 | 1.95 | 0.027** | 2.35 |
| 113 | 0.33 | 0.372 | 0.60 | -0.08 | 0.531 | -0.11 | 1.38 | 0.086* | 1.62 |
| 114 | -0.05 | 0.520 | -0.10 | 0.91 | 0.181 | 1.38 | 0.68 | 0.249 | 0.90 |
| 115 | 0.17 | 0.434 | 0.42 | 0.65 | 0.258 | 0.89 | 1.43 | 0.079* | 1.94 |
| 116 | 1.27 | 0.106 | 2.60 | 0.49 | 0.313 | 0.73 | 2.18 | 0.016** | 2.58 |
| 117 | 1.75 | 0.043** | 3.41 | 0.85 | 0.200 | 1.12 | 2.30 | 0.012** | 2.48 |
| 118 | 1.90 | 0.031** | 3.27 | 0.87 | 0.193 | 1.07 | 1.89 | 0.031** | 2.41 |
| 119 | 1.12 | 0.134 | 2.36 | 0.67 | 0.252 | 0.83 | 2.03 | 0.023** | 2.73 |
| 120 | 1.25 | 0.108 | 2.97 | -0.05 | 0.518 | -0.06 | 0.55 | 0.292 | 0.78 |
| 121 | 1.37 | 0.089* | 3.14 | 0.17 | 0.434 | 0.21 | 0.89 | 0.187 | 1.21 |
| 122 | 2.25 | 0.014** | 4.70 | 0.18 | 0.428 | 0.24 | 0.22 | 0.415 | 0.28 |
| 123 | 1.88 | 0.032** | 4.04 | 0.69 | 0.245 | 0.96 | 0.49 | 0.314 | 0.67 |
| 124 | 2.96 | 0.002** | 6.75 | 1.25 | 0.108 | 1.74 | 0.53 | 0.300 | 0.82 |
| 125 | 2.25 | 0.014** | 5.32 | 1.85 | 0.034** | 2.61 | 0.24 | 0.407 | 0.33 |
| 126 | 1.82 | 0.037** | 4.15 | 1.99 | 0.025** | 2.87 | -0.21 | 0.583 | -0.33 |
| 127 | 1.77 | 0.041** | 4.21 | 0.97 | 0.168 | 1.52 | 0.55 | 0.291 | 0.83 |
| 128 | 1.77 | 0.041** | 3.22 | 1.59 | 0.057* | 2.24 | 1.60 | 0.057* | 2.14 |
| 129 | 1.68 | 0.049** | 3.37 | 0.82 | 0.207 | 1.14 | 1.36 | 0.089* | 1.89 |
| 130 | 1.37 | 0.089* | 2.65 | 0.47 | 0.318 | 0.66 | 0.23 | 0.409 | 0.36 |
| 131 | 1.30 | 0.099* | 2.49 | -0.32 | 0.625 | -0.50 | 0.49 | 0.311 | 0.67 |
| 132 | 0.67 | 0.254 | 0.99 | -0.14 | 0.556 | -0.22 | 0.66 | 0.257 | 0.80 |
| 133 | 0.78 | 0.220 | 1.53 | -0.18 | 0.570 | -0.23 | 0.25 | 0.402 | 0.31 |
| 134 | 0.84 | 0.203 | 1.68 | 1.01 | 0.158 | 1.40 | 0.01 | 0.496 | 0.01 |
| 135 | 2.78 | 0.004** | 6.12 | 1.65 | 0.051* | 2.11 | 0.62 | 0.267 | 0.86 |
| 136 | 2.58 | 0.006** | 6.10 | 0.64 | 0.263 | 0.82 | 1.21 | 0.115 | 1.71 |
| 137 | 3.02 | 0.002** | 5.58 | 1.75 | 0.042** | 2.24 | 1.29 | 0.101 | 1.86 |
| 138 | 1.95 | 0.028** | 3.76 | 1.09 | 0.138 | 1.46 | 1.34 | 0.091 | 2.05 |
| 139 | 1.94 | 0.029** | 4.63 | 0.22 | 0.412 | 0.27 | 1.08 | 0.142 | 1.65 |
| 140 | 1.40 | 0.084* | 3.58 | -0.68 | 0.750 | -0.89 | 0.64 | 0.262 | 1.09 |

Note: * *p* < .10, ** *p* < .05, *** *p* < .001
